# Pangenome calculation beyond the species level using RIBAP: A comprehensive bacterial core genome annotation pipeline based on Roary and pairwise ILPs

**DOI:** 10.1101/2023.05.05.539552

**Authors:** Kevin Lamkiewicz, Lisa-Marie Barf, Konrad Sachse, Martin Hölzer

## Abstract

Pangenome analysis is a computational method for identifying genes that are present or absent from a group of genomes, which helps to understand evolutionary relationships and to identify essential genes. While current state-of-the-art approaches for calculating pangenomes comprise various software tools and algorithms, these methods can have limitations such as low sensitivity, specificity, and poor performance on specific genome compositions. A common task is the identification of core genes, i.e., genes that are present in (almost) all input genomes. However, especially for species with high sequence diversity, e.g., higher taxonomic orders like genera or families, identifying core genes is challenging for current methods. We developed RIBAP (Roary ILP Bacterial core Annotation Pipeline) to specifically address these limitations. RIBAP utilizes an integer linear programming (ILP) approach that refines the gene clusters initially predicted by the pangenome pipeline Roary. Our approach performs pairwise all-versus-all sequence similarity searches on all annotated genes for the input genomes and translates the results into an ILP formulation. With the help of these ILPs, RIBAP has successfully handled the complexity and diversity of *Chlamydia, Klebsiella, Brucella, and Enterococcus* genomes, even when genomes of different species are part of the analysis. We compared the results of RIBAP with other established and recent pangenome tools (Roary, Panaroo, PPanGGOLiN) and showed that RIBAP identifies all-encompassing core gene sets, especially at the genus level. RIBAP is freely available as a Nextflow pipeline under the GPL3 license: https://github.com/hoelzer-lab/ribap.

## Introduction

Pangenomics aims to understand the whole genomic content of a species or population, including both the core genome (conserved genes shared by all or nearly all members of the group) and the accessory genome (variable genes that are present in only a subset of members) (Mira et al. 2010; Tettelin et al. 2005). Besides “core genome” and “accessory genome”, terms like “persistent”, “shell”, and “cloud” are used to describe different sets of genes based on their presence and distribution across a given set of genomes. In general, determining the pangenome allows for comparing multiple genes and identifying evolutionary relationships (3), thus providing new insights into bacterial pathogenicity and clinical microbiology (Anani et al. 2020). Overall, classifying genes into categories such as “core” and “accessory” allows insights into the evolution and adaptation of a particular species or group of species and helps researchers identify critical functional genes that may be important for understanding the biology and ecology of these organisms. This can be particularly important in bacteria because of their high genetic diversity and ability to exchange genetic material through mechanisms such as horizontal gene transfer.

Computational pangenome analysis pipelines are used to calculate a pangenome by comparing the genomes (genes) of multiple individuals within a species or population (for an overview, see (Pantoja et al. 2020; Bonnici, Maresi, and Giugno 2021)). Recent tools, such as Roary (Page et al. 2015), Panaroo (Tonkin-Hill et al. 2020), and PPanGGOLiN (Gautreau et al. 2020), typically involve aligning the genomes/genes and identifying shared and unique genes. To this end, the genes are annotated using tools such as Prokka (Seemann 2014) or Bakta (Schwengers et al. 2021) before being provided as input for gene-oriented approaches to pangenome content discovery.

One challenge in pangenomics is dealing with a large amount of data generated by analyzing multiple genomes. This can require significant computational resources and expertise in bioinformatics. Additionally, accurately predicting the functions of genes can be difficult, especially for those present in only a small subset of individuals. Finally, pangenomics studies often involve analyzing diverse populations, which can be difficult to define and sample accurately. In this context, a particular problem arises when sequence similarity between genes belonging to the core genome is low, for example, when calculating a pangenome for diverse species beyond the genus level. In this case, it may be difficult to correctly assign genes to the core genome, and they may erroneously end up as independent groups in the accessory genome. Thus, defining homology based on sequence similarity alone often underestimates the true core genome, especially when comparing genomes across species or genus boundaries. Simply scaling up established bioinformatics pipelines will not be sufficient to realize the full potential of rapidly growing and diverse genomic datasets (The Computational Pan-Genomics Consortium 2018). Therefore, new, qualitatively different computational methods and paradigms are needed to advance the field of computational pangenomics.

Here we present RIBAP (Roary ILP Bacterial core Annotation Pipeline), a comprehensive bacterial pangenome annotation pipeline based on Roary (Page et al. 2015) and pairwise integer linear programs (ILPs) as introduced by (Martinez et al. 2015). The development of the pipeline was motivated by our comparative genome studies on different *Chlamydia* species (Hölzer et al. 2020; Vorimore et al. 2021; Sachse et al. 2022). Here, we could not calculate meaningful core genome sets based on experts’ evaluation using the available pangenome tools without lowering the sequence similarity cutoff well below the values recommended by the pipeline authors. Therefore, we decided to keep the initial pangenome calculations with high thresholds for sequence similarity and refine the resulting robust gene sets as we proceeded, ending in implementing the RIBAP pipeline. RIBAP combines robust sequence homology information from Roary with pairwise ILP calculations to produce a complete core gene set - even at the genus level. First, RIBAP performs annotations with Prokka (Seemann 2014), calculates a pangenome with Roary, refined by pairwise ILPs, and finally visualizes the results in an interactive HTML table linking each gene family of the pangenome to its multiple sequence alignment and sequence-based phylogenetic tree. RIBAP is implemented in Nextflow (Di Tommaso et al. 2017) and comes with Docker/Singularity/Conda support for easy installation and execution on local machines, high-performance clusters, or the cloud.

## Material & Methods

### Used bacterial datasets in this study

We selected four bacterial datasets with different compositions to evaluate the performance of RIBAP: *Enterococcus* (44 genomes), *Brucella* (71), *Chlamydia* (102), and *Klebsiella* (167). We selected *Enterococcus* as a representative of gram-positive bacteria ubiquitous in various environmental settings and with a diverse genome size range from 2.6 to 4.2 Mbp. The *Enterococcus* dataset is composed of the species *E. faecium* (21 genomes), *E. faecalis* (14), *E. durans* (2), *E. hirae* (2), *E. casseliflavus* (1), *E. gallinarum* (1), *E. mundtii* (1), *E. silesiacus* (1), and *E. sp*. (1). *Brucella* are animal pathogenic, Gram-negative bacteria. Our dataset includes genomes ranging in size from 3.2 - 3.6 Mbp with the species *B. melitensis* (24), *B. suis* (16), *B. abortus* (14), *B. canis* (6), *B. sp*. (4), *B. pinnipedialis* (2), *B. ceti* (2), *B. microti* (1), *B. ovis* (1), and *B. vulpis* (1). The *Chlamydia* dataset, gram-negative and human and animal pathogenic bacteria, contains the dataset with the smallest genomes in the range of 1 - 1.2 Mbp and includes the species of *C. trachomatis* (70), *C. psittaci* (15), *C. muridarum* (5), *C. pecorum* (3), *C. abortus* (3), *C. gallinacea* (2), *C. avium* (1), *C. felis* (1), *C. pneumoniae* (1), and *C. suis* (1). Finally, our largest dataset consists of *Klebsiella* species, gram-negative and human pathogenic bacteria with the largest genome sizes in our benchmark of 5.1 - 7.3 Mbp. The species included are *K. pneumoniae* (134), *K. oxytoca* (8), *K. variicola* (7), *K. aerogenes* (6), *K. michiganensis* (6), *K. quasipneumoniae* (4), and *K. sp*. (2). All genomes were downloaded from NCBI and are also available here: https://osf.io/g52rb; their accession IDs are summarized in Supplementary Table S1. For each described genus, we further selected the species with the most genomes to assess the performance of RIBAP. For each dataset, we calculated the percentage of conserved proteins (POCP, https://github.com/hoelzer/pocp v1.1.1) (Qin et al. 2014) to examine how similar the selected genomes are at the protein level.

### General workflow of RIBAP

The RIBAP pipeline (Figure 1) is implemented in Nextflow, a workflow management system for reproducible analyses (Di Tommaso et al. 2017). Each tool dependency is solved via Conda environments or pre-build Docker/Singularity containers (Boettiger 2015). To ensure compatibility between genome annotations, the pipeline begins by (re-)annotating all input genomes with Prokka, a popular tool that identifies bacterial gene features such as protein-coding sequences (CDS), tRNAs, and rRNAs (Seemann 2014). These annotations are then used to perform pairwise all-versus-all sequence similarity searches with MMSeqs2 (Steinegger and Söding 2017). The results of these searches are used to generate ILP problems, which are subsequently solved with GLPK (Free Software Foundation 2020). In addition to the MMSeqs2 analyses, the pipeline also uses Roary (Page et al. 2015) to calculate a pangenome scaffold, which is refined with the help of the ILPs (see the section below for details). The final step of the pipeline is to link and potentially expand homologous gene families in the Roary scaffold (called “Roary clusters”) using the individual results of the ILP analyses into so-called “RIBAP groups” (Figure 2). We consider every gene that is present in all input genomes as a core gene. For each RIBAP group, we calculate a multiple sequence alignment (MSA) and a phylogenetic tree with MAFFT (Katoh and Standley 2013) and FastTree (Price, Dehal, and Arkin 2009), respectively. Optionally, the user can further calculate a phylogenetic tree based on the complete core gene set using IQ-TREE 2 (Minh et al. 2020). To reduce runtime, we apply CD-HIT (W. Li and Godzik 2006) on each core gene set MSA and remove MSAs from the core gene set phylogeny calculation that lack diversity. RIBAP summarizes the results in an interactive HTML file, providing a searchable table and access to all alignments and phylogenetic trees for each gene family. All tool versions and the detailed descriptions of the individual steps are based on the release version 1.0.0 of RIBAP (https://github.com/hoelzer-lab/ribap).

**Figure 1:**
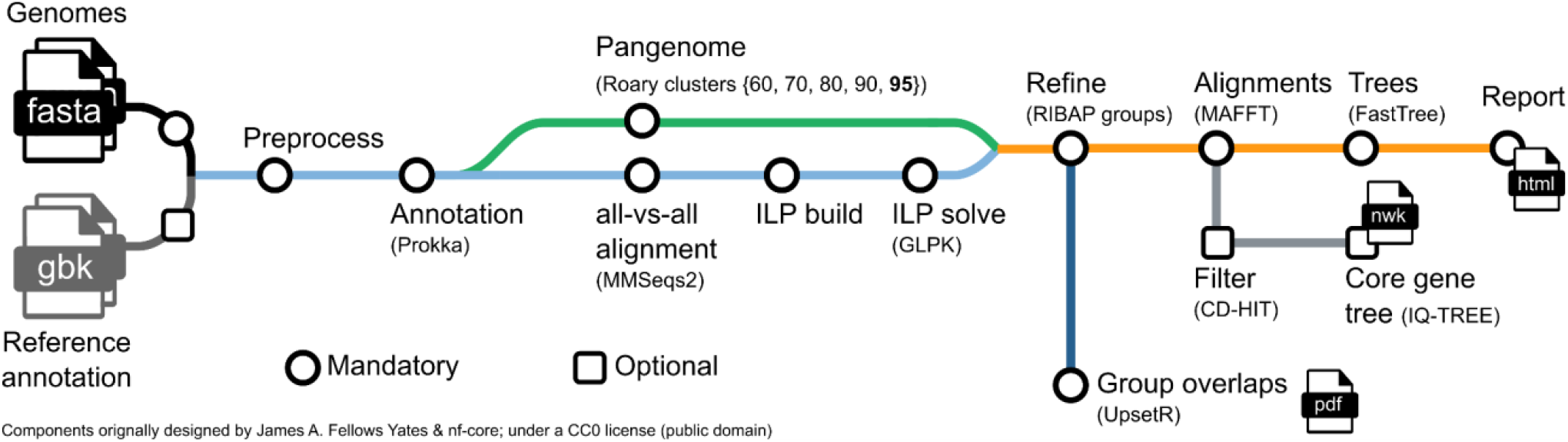
Schematic overview of the RIBAP pipeline. The only mandatory input are genomes in FASTA format that can be provided directly or via a CSV file of the paths. Reference annotations in GenBank format (gbk) can be provided as optional input to guide Prokka gene annotations. The pipeline will calculate a scaffold pangenome producing Roary gene clusters, which are further refined by the ILP results into so-called RIBAP groups. For the genes within each RIBAP group, a multiple sequence alignment (MSA) and a phylogenetic tree are calculated and linked in the final summary report table in HTML format. Optionally, a tree (Newick format, nwk) for the concatenated core gene MSAs can be calculated. We use CD-HIT, to remove MSAs that are only composed of identical sequences before tree calculation. An UpSet plot visually summarizes overlaps between the identified RIBAP groups of all analyzed genomes. All intermediate output files are provided for detailed investigation and further downstream analyses.

**Figure 2:**
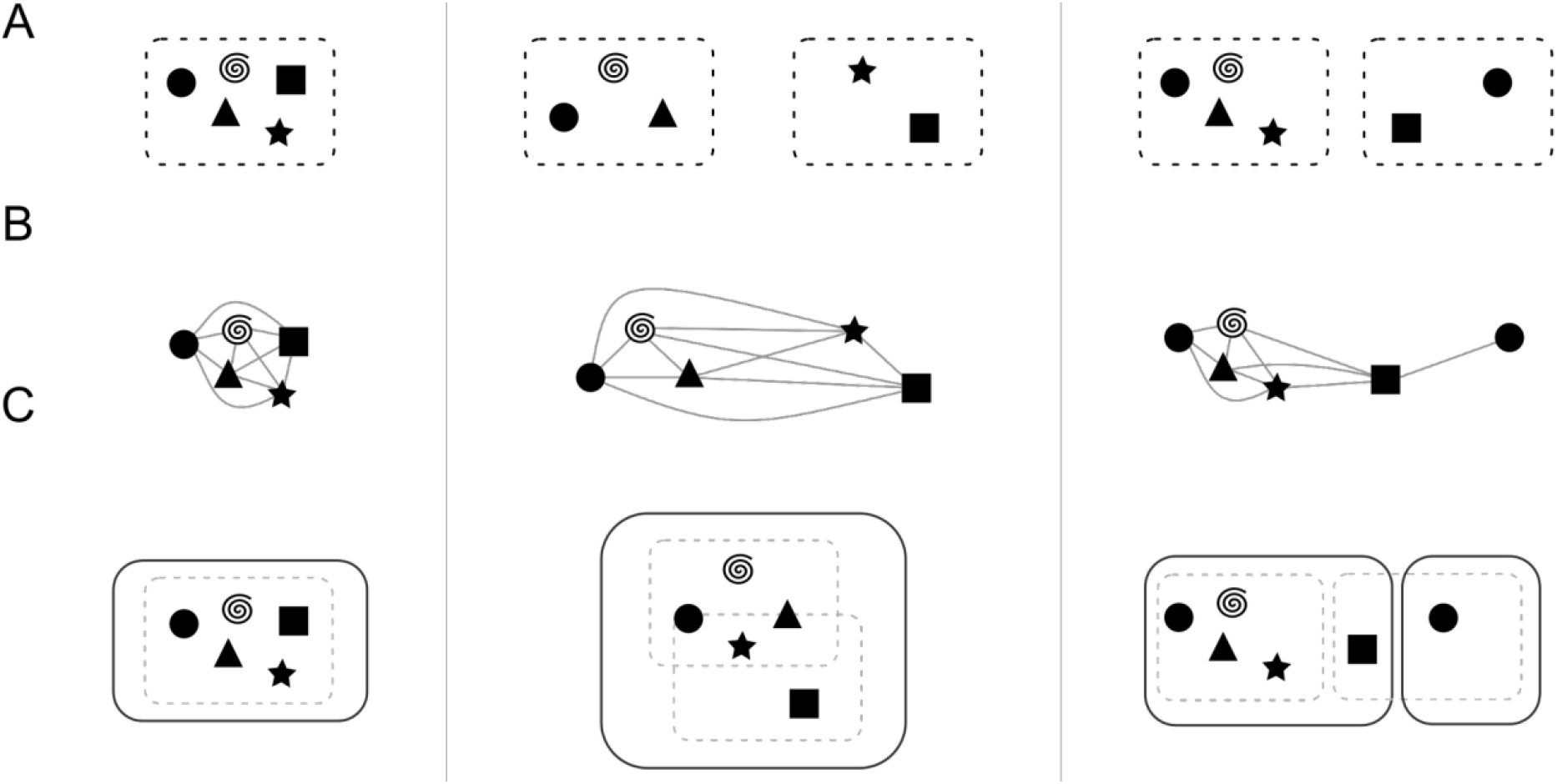
General combination scheme of the Roary and ILP results. The left-hand side describes a trivial case, showing a Roary cluster with five genes that is also a RIBAP group. The middle panel shows two Roary clusters (three and two genes, respectively) that are finally merged into one RIBAP group with the help of the ILPs. The right-hand panel shows again two Roary clusters that result in two RIBAP groups. The smaller RIBAP group is labeled as a subgroup of the larger RIBAP group. **(A)** The original Roary clusters as determined at a sequence similarity threshold of 95%. **(B)** Extracted genes and their pairwise ILP connections. **(C)** The resulting RIBAP groups (and the original Roary clusters) after our merging procedure.

### Initial gene annotation

RIBAP utilizes Prokka (Seemann 2014) (v1.14.6, default parameters) to annotate all input genomes. Each CDS, defined from start to stop codon, is searched in a protein database derived from UniProtKB. Coding regions without a database hit are labeled as “hypothetical protein” by Prokka. In addition, Prokka annotates rRNA and tRNA genes. While the gene annotation itself does not affect the calculations of RIBAP, the genomic coordinates of each CDS are used to perform subsequent steps in our pipeline. The annotation itself is again included when results are summarized in the tabular output. Providing a reference annotation file in GenBank format to guide the Prokka annotations is also possible. A CSV file can be provided to guide the genome annotation using different reference annotations.

### Roary pangenome calculation

Based on the Prokka annotations, we calculate a preliminary scaffold pangenome using the tool Roary (Page et al. 2015) (v3.13.0, default parameters except for sequence similarity thresholds). Roary outputs homologous genes potentially belonging to a group into clusters. The threshold for sequence similarity is set to 95% by default, and the corresponding results (Roary clusters) are used for subsequent analysis steps, e.g., for merging with the ILP results. However, RIBAP performs additional Roary calculations with lower thresholds (60%, 70%, 80%, 90%). The Roary clusters resulting from these lower similarity thresholds are not used for downstream calculations but for visualization and comparison.

### Pangenome refinement via Integer Linear Programming

We refine the initial Roary clusters based on 95% sequence similarity to tackle the issue of common pangenome calculation tools of underestimating the number of core genes in genomes with high sequence diversity or in the context of inconsistent gene annotations (T. Li and Yin 2022; Zhou, Charlesworth, and Achtman 2020). Our approach utilizes individual, pairwise comparisons of the genes of all input genomes and refines the scaffold pangenome as calculated by Roary. First, all gene features, as predicted by Prokka (mainly CDS, but also tRNAs and rRNAs), are used in an MMSeqs2 (Steinegger and Söding 2017) (v10.6d92c) all-vs-all comparison. We split this output into all possible pairwise comparisons between the input genomes and use these pairwise comparisons to formulate ILPs, which we subsequently solve using the GNU Linear Programming Kit (GLPK, v4.65) package (Free Software Foundation 2020). To limit the run-time of RIBAP, per default, each ILP has a time limit of 240 seconds (--tmlim 240s in GLPK). For a brief overview of our ILP formulation, check below.

To further reduce the run-time of RIBAP, we split the ILP problem of two genomes into several sub-ILPs based on disjoint components in the initial adjacency graph. Further, trivial cases where a direct one-to-one mapping of genes is possible are not parsed into an ILP problem but are directly accepted as homologs by our ILP approach.

The ILPs provide homology mappings between genes of lower sequence similarity (60% or higher). Thus, we have a scaffold pangenome calculated by Roary and all pairwise sets of homologous genes given any two input genomes. This information is merged in the following fashion (visualized in Figure 2): First, we extract all genes for each Roary cluster identified using a 95% similarity threshold. Then, we compare hits of each gene in our pairwise ILPs with the information Roary provided. In the trivial case, no new information is added with the inclusion of our ILPs (see Figure 2, left). However, if any homolog gene derived from the ILPs belongs to a different Roary cluster, the two clusters are merged into a preliminary RIBAP group (Figure 2, middle and right). To account for gene duplications (i.e., paralogs), we further refine a RIBAP group. If any genome has two or more genes within the same RIBAP group, we define subgroups for each paralog gene in the original preliminary RIBAP group (Figure 2, right). Let *g_A_* be a set of genes that are all paralogs in a genome *A*. To resolve the issue of determining and selecting the “main” homolog gene for all other genomes within this RIBAP group, we compare the individual ILP scores and the Roary score. First, we evaluate the number of hits based on our pairwise ILPs, i.e., if one gene is connected to the rest of the cluster more often than the other gene, we pick this as the “main” homolog. If this is ambiguous, we fall back to the scaffold pangenome determined by Roary. For each gene in *g_A_*, we check the cluster sizes these genes belong to and determine the gene with the largest cluster to be the “main” homolog. If this second analysis still yields ambiguity, we make the best guess based on the Prokka annotation and gene name. The rest of *g_A_* is then split into *n*−1 subgroups, where *n* is the size of *g_A_*. If there are two or more genomes with paralogs, we repeat the procedure for each subgroup.

### FFDCJ distance and ILP formulation

To refine the pangenome calculation by Roary, we employ all-vs-all comparisons of annotated genes for each pair of genomes. Let *A* and *B* be such a pair of genomes with *n* and *m* genes, respectively. We use the annotation of Prokka to determine *n* and *m*, but we do not use the functional annotation, the gene names, itself, to determine further homology. Each gene *A_i_* with *i ϵ*{1..*n*} is compared with each gene *B_j_* with *j ϵ*{1..*m*}. This leads to sequence similarities (and potential orientation differences) between each pair of genes of the two genomes. Following previous studies, we first construct a gene similarity graph *GS_σ_(A,B)* (Figure 3A) based on the two genomes *A* and *B* and all gene similarities encoded by *σ* (Braga et al. 2013). We use the reported bitscore of MMseqs2 as a combined value representative for the sequence similarity and alignment length of two genes. Now, let *M* be a matching of *GS_σ_(A,B)*, then *A^M^* and *B^M^* denote the reduced genomes of *A* and *B*. In reduced genomes, singletons derived from indel events are removed (Figure 3B). Due to the orientation of a gene, we can distinguish the two ends of a gene called extremities (t – gene tail, or the 3’ end; h – gene head, or the 5’ end). We now build the adjacency graph *AG_σ_*(*A^M^,B^M^*) by modeling a gene’s adjacency via the two neighboring genes’ extremities (Figure 3C). Here, assuming identical genome organization, *AG_σ_*(*A^M^,B^M^*) would result in cycles of length two (adjacent genes). We refer to these two elements as “fixed components” as no genome rearrangement events are needed to transfer one genome to another. For all other graphs, genome rearrangements have to be applied. On the other hand, we have to consider gene similarities if we calculate the distance between two genomes without gene family assignments. Sequence similarities and genomic organization between *A* and *B* could be contradictory, e.g., depending on whether one prefers (slightly) higher individual similarities or fewer genomic rearrangements such as inversions or transpositions. This contradiction leads to an optimization problem described by Martinez *et al*., named the family-free DCJ (FFDCJ) distance (Martinez et al. 2015). Martinez *et al*. proposed an ILP to compute the optimal FFDCJ distance between two genomes *A* and *B*. This optimal FFDCJ distance of two genomes *A* and *B* is defined as given in Equation 1, where |*M*| is the size of the maximum matching in *GS_σ_(A,B),c*, is the number of cycles in *AG_σ_*(*A^M^,B^M^*) and *ω(M)* are the summed weights of the edges in the matching. The parameter *α*ϵ{0,1} weights the genome order and the sum of individual gene similarities.

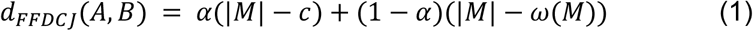

We extended this ILP formulation to consider indel events (as depicted in Figure 3B) (Braga, Willing, and Stoye 2011). First, we label each gene not part of a fixed component as a potential indel event. Next, we summarize consecutive indel events into a block (Braga, Willing, and Stoye 2011; Braga et al. 2011). This is motivated by the fact that it seems reasonable to have larger indel events, affecting consecutive genes at once, instead of having many individual indel events. Similarly to how Martinez *et al*. count cycles (see (Martinez et al. 2015; Shao, Lin, and Moret 2015)), we count blocks of indels and consider them in the objective function. For this, two adaptations of the original ILP have been made: (i) for each singleton, let there be an edge in *GS_σ_(A,B)* that connects the two gene extremities of the singleton in its genome. We call this edge a self-edge (see (Martinez et al. 2015)) and include its cost to the objective function of the ILP. We (ii) define a binary variable *b_i_* that indicates whether a gene *i* is at the end of a block (Braga et al. 2011). The number of blocks (i.e. number of *b_i_* set to 1) is also included in the objective function. The weights of self edges and blocks are determined by *α* (default: 0.5). These adaptations lead to our (naive) FFDCJ-indel distance as given in Equation 2. It extends Equation 1 by adding the number of singletons *S* and number of indel blocks *I* to the rearrangement part of the equation. Note that we are still looking for maximum matchings, similar to the original ILP. Therefore, an indel event is only considered if there is no way to match a gene to the other genome. Additionally, we penalize indel events twice by our adaptation; once for every singleton and another time for each block of indels. This is based on our observations that only considering one of the two adaptations led to a dramatic overestimation of indels (adaptation (i)) or of the ILP interpreting genome *A* as one deletion block and genome *B* as one insertion block (adaptation (ii)).

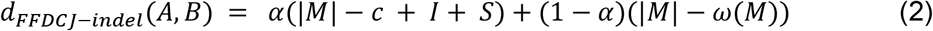

**Figure 3:**
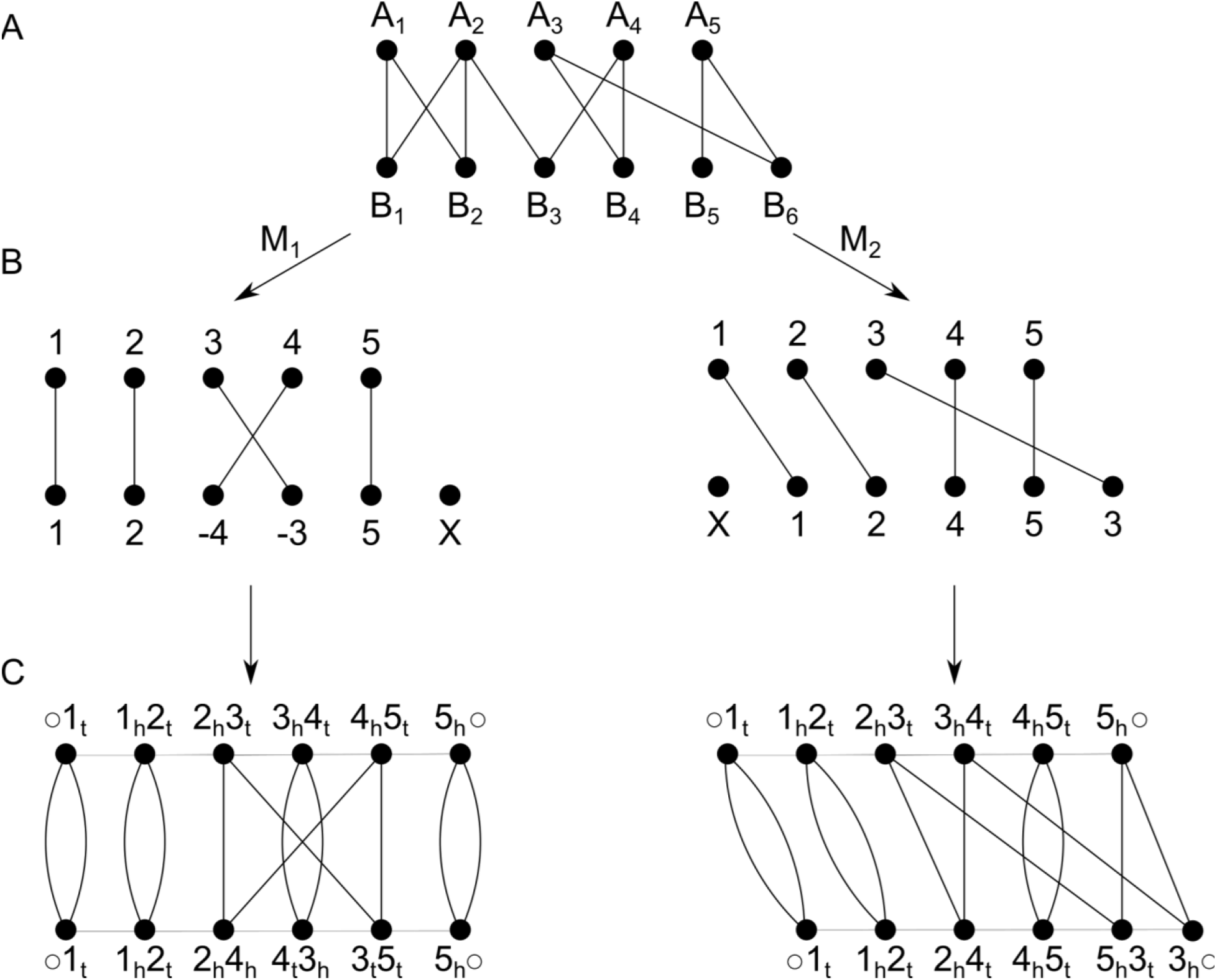
**(A)** Gene similarity graph of two genomes, A and B, with five and six genes, respectively. Note that, for simplicity, edge weights are omitted in this figure. **(B)** Two possible matchings of the graph. Both contain an indel event. Additionally, M1 contains an inversion and M2 a transposition. **(C)** Derived adjacency graph of the two matchings. Each gene is denoted by its gene extremities, black edges denote homology across A and B, gray edges represent an adjacency within a genome. t – gene tail, or the 3’ end; h – gene head, or the 5’ end.

### Alignment, tree, and summary output

For each RIBAP group, we calculate a multiple sequence alignment (MSA) and a phylogenetic tree with MAFFT (v7.455, default parameters) (Katoh and Standley 2013) and FastTree (2.1.10, default parameters) (Price, Dehal, and Arkin 2009), respectively. Lastly, we produce an interactive HTML file, which visualizes the pipeline results in a searchable table and links to each MSA and tree. To visualize the pangenome, we employ an UpSet plot with the UpSetR package (v1.4.0) (Conway, Lex, and Gehlenborg 2017). The user can also activate the calculation of a phylogenetic tree based on all core gene MSAs using IQ-TREE 2 (Minh et al. 2020) (v2.2.0.3, -spp mode). To reduce runtime, we apply CD-HIT (W. Li and Godzik 2006) on each core gene set MSA and remove MSAs from the core gene set phylogeny calculation that are composed of 100% identical sequences. The remaining MSAs are then individually processed by IQ-TREE 2 to estimate the best-fitting model for each gene, respectively.

## Results & Discussion

Based on rapid advances in sequencing technologies and computational approaches in the past two decades, the classification of bacterial genes into homologous groups based on their presence or absence has become a common comparative task referred to as microbial pangenomics (Mira et al. 2010; Gmiter et al. 2021; Medini et al. 2005). Today, researchers have access to various tools (Vernikos 2020) to input genomes or genes to define accessory and core genes or, at an even finer granularity, shell, cloud, and persistent genes. The resulting gene groupings depend on the applied computational tool (Pantoja et al. 2020; Bonnici, Maresi, and Giugno 2021; The Computational Pan-Genomics Consortium 2018) and parameter settings, e.g., sequence similarity thresholds or the relative number of input genomes required to make up a specific group. In our experience, the composition of the input genomes and their sequence similarity reflecting their evolutionary relatedness can be challenging for computational pangenome tools, leading to underestimation of core gene sets or false positive assignments due to too low sequence similarity thresholds. Therefore, we developed RIBAP to refine the robust gene clusters computed by Roary based on high sequence similarity by combining it with an ILP approach for calculating core gene groups, especially for evolutionarily diverse genome inputs.

### RIBAP reconstructs more comprehensive core genomes when dealing with diverse input genomes

To compare the performance of RIBAP, we analyzed the results of different tools commonly used to calculate pangenomes. While such tools perform well on input genomes from the same taxonomic species, core genomes are underestimated when sequence diversity increases. Thus, we applied RIBAP and three other tools (Roary, Panaroo, PPanGGOLiN) to different bacterial datasets, as summarized in Table 1 and Supplementary Table S1. To challenge the tools, we deliberately chose the datasets so that genomes with lower sequence similarity were also included. To investigate how similar the selected genomes are at the protein level, we calculated pairwise POCP (percentage of conserved proteins) values for the genomes belonging to each species (Qin et al. 2014). POCP is a metric to evaluate the similarity between two bacterial genomes. It is also a widely accepted parameter for delimiting genus boundaries in prokaryotic genome-based taxonomy (Qin et al. 2014). The POCP is calculated by comparing the number of orthologous proteins shared between two bacterial genomes and dividing it by the total number of proteins in both genomes. The POCP values showed that each of our datasets includes highly similar and more distant genomes (see https://osf.io/g52rb and Supplementary Table S2). For example, the *Brucella* dataset includes three genomes with POCP values ∼89% (09RB8471, 09RB8910, 141012304), while most genomes have a POCP significantly larger 90%. The POCP calculation also highlights *Brucella vulpis* strain F60 with POCPs ∼91% as more distant in this dataset. Another extreme example is *Klebsiella michiganensis* strain RC10, which has a POCP of only 60% compared to *Klebsiella oxytoca* strain CAV1374 and a generally low POCP in this dataset.

**Table 1.** Core genome sizes of selected bacterial datasets as predicted by different tools used in this study. Sizes were extracted from the gene absence/presence tables of respective tools. The number of total genes was calculated based on the average number of Prokka gene annotations. The average POCP values were calculated from Supplementary Table S2.

| Dataset | # input genomes | RIBAP | Roary (95%) | Panaroo | PPanGGOLiN | Average genes annotated | Average POCP value [%] |
| --- | --- | --- | --- | --- | --- | --- | --- |
| *Brucella* spp. | 71 | 2,444 | 1,602 | 2,342 | 2,121 | 3,151.66 | 97.22 |
| *Brucella melitensis* | 24 | 3,037 | 2,949 | 3,086 | 2,955 | 3,153.17 | 97.39 |
| *Chlamydia* spp. | 102 | 772 | 9 | 0 | 127 | 922.46 | 91.85 |
| *Chlamydia trachomatis* | 71 | 866 | 826 | 868 | 853 | 989.31 | 98.97 |
| *Enterococcus* spp. | 44 | 1,491 | 21 | 55 | 346 | 2,986.95 | 69.60 |
| *Enterococcus faecium* | 21 | 1,989 | 1,409 | 1,887 | 1,850 | 3,001.67 | 89.40 |
| *Klebsiella* spp. | 167 | 3,205 | 923 | 2,690 | 1,581 | 5,347.36 | 86.44 |
| *Klebsiella pneumoniae* | 134 | 3,748 | 2,974 | 3,694 | 3,278 | 5,298.09 | 89.55 |

All evaluated tools generally provide a similar core genome size when input genomes are taken from the same species. However, including genomes from different species of the same genus decreases the size of core genomes for all bacterial genera tested. Most surprising are the results for *Chlamydia* and *Enterococcus*. Here, RIBAP is the only tool calculating plausible core genome sizes. While other tools compute core gene sets containing only 0 - 13.77% (*Chlamydia* spp.) and 0.7 - 11.58% (*Enterococcus* spp.) of all annotated genes (based on Table 1, gene counts averaged over all genomes in the respective dataset), RIBAP’s core gene set covers 83.69% and 49.92%, respectively. These results are also more consistent with previously published core gene sets calculated on less diverse input datasets of *Chlamydia* spp. and *Enterococcus* spp. (Khan, Jalal, and Uddin 2023; Sigalova et al. 2019). In general, when sequence identity among CDS is low, pangenome tools are challenged to identify homologous genes. This is further indicated by POCP calculations of the datasets.

As mentioned above, our POCP analysis of *Brucella* spp. genomes revealed high inter-species genome similarity (Table 1). This is also indicated by the performance of all pangenome calculation tools, as the *Brucella* dataset is the most robust one in our assessment when genomes from different species are included as input. Given the high POCP values of this dataset (*B. melitensis* 97.39% and *Brucella* spp. 97.22%), we noticed that the genomes selected of this bacterial genus are more conserved than others and thus facilitate core genome calculation based on sequence similarity thresholds. This may be a consequence of the (historic) taxonomic classification of brucella strains, which is characterized by relatively high sequence similarity thresholds (Whatmore 2009; Ficht 2010).

We did not observe a drastic drop in core genome size for the *Klebsiella* dataset. As indicated by our POCP analysis, the selected genomes representing the *Klebsiella* genus had a relatively high pairwise sequence similarity, causing tools with strict sequence similarity thresholds to still recover many core genes. While one outlier (*Klebsiella michiganensis* strain RC10) had POCP values of around 65%, the remaining *Klebsiella* genomes had an average value of 86.44%. The corresponding value of *K. pneumoniae* was only slightly higher at 89.55%. Still, compared to the other tools, RIBAP recovered the largest core genome of *Klebsiella* spp. (around 60% of the annotated genes, Table 1). Roary, Panaroo, and PPanGGoLiN predicted the core genome size of *Klebsiella* spp. to be around 17.26%, 50.30%, and 29.57% of the annotated genes, respectively. Further, comparing the *Klebsiella* spp. core genome sizes with the predicted core genome sizes of the *K. pneumoniae* dataset supports our hypothesis that other tools are challenged by diverse input genomes. Relative to the predicted size of the *K. pneumoniae* core genome, RIBAP recovered 85.5% (3,205 of 3,748) of core genes in the *Klebsiella* spp. dataset, while Roary, Panaroo, and PPanGGoLiN only recovered 31.03% (923 of 2,974), 72.82% (2,690 of 3,694), and 48.23% (1,581 of 3,278), respectively. A small reduction in POCP values thus caused tools to lose many core genes.

In contrast, the *Chlamydia* dataset comprising the entire genus challenged state-of-the-art tools. POCP values ranged between ∼75% and above 99% for this dataset, where *C. pneumoniae* had the lowest values on average. Considering only *C. trachomatis*, POCP values are above 98% for each pairwise comparison, leading to sound core genomes for this species. However, including other species with lower POCP values causes core genome sizes to decrease dramatically. While each tool calculates over 800 genes to be part of the core genome for *C. trachomatis*, the core genomes for the *Chlamydia* spp. are reduced to 9 (Roary), 0 (Panaroo), and 127 (PPanGGOLiN) genes, respectively. Only RIBAP calculates a core genome with a reasonable size of 772 genes. Recent studies estimate the core genome size to be around 880 (*C. trachomatis*) and 700 (*Chlamydia* spp.) genes, respectively (Versteeg et al. 2018; Sigalova et al. 2019).

We made similar observations with the *Enterococcus* dataset. Here, the species *E. faecium* has a pairwise POCP value between ∼81% and 99%, leading to similar core genome sizes with different tools. However, including genomes from *Enterococcus* spp. resulted in POCP values as low as ∼43%. As expected, core genome size decreased from around 1,900 to 21 (Roary), 55 (Panaroo), and 346 (PPanGGOLiN) genes, respectively. Our refinement approach, including the ILPs, resolved many Roary clusters and proposed a core genome size of 1,491 genes. Thus, the core genome size of RIBAP for *Enterococcus* spp. covers 74.96% (1,491 of 1,989) of the core genome size of *E. faecium* at the species level, while Roary (1.49%), Panaroo (2.91%), and PPanGGOLiN (18.70%) calculate much smaller core genome sets at the genus level compared to the respective species level (Table 1).

### RIBAP identifies core genes with low sequence similarity from diverse input genomes

To emphasize the advantages of RIBAP, we looked at the *ompA* gene, which is present in all species of *Chlamydia*. This gene encodes the major outer membrane protein or porin, which researchers have been using to subdivide the major species of *Chlamydia* into different serotypes based on recognized epitopes on the protein surface (Stephens et al. 1982; Wang et al. 1985). As shown in Figure 4, the protein sequence similarity of OmpA in different species of *Chlamydia* can be as low as around 60%. Due to the ILP refinement implemented in RIBAP, we can reconstruct this core gene despite its high sequence diversity. In contrast, Roary, Panaroo, and PPanGGOLiN do not detect *ompA* as a core gene of *Chlamydia* spp. when used with default parameters. Further, using the default parameter of Roary (sequence similarity threshold of 95%), Figure 4 also indicates that Roary would not even detect *ompA* as a core gene for the individual species *C. trachomatis* or *C. psittaci*, respectively. In both cases, sequence similarity would have to be reduced to 80% to recognize *ompA* as a core gene with Roary (Figure 4). This further supports our point that many pangenome calculation tools underestimate the actual number of core genes, even if genomes from the same species are used as input. In this context, please note that RIBAP currently defines a gene as part of the core genome if it is present in all input genomes.

**Figure 4:**
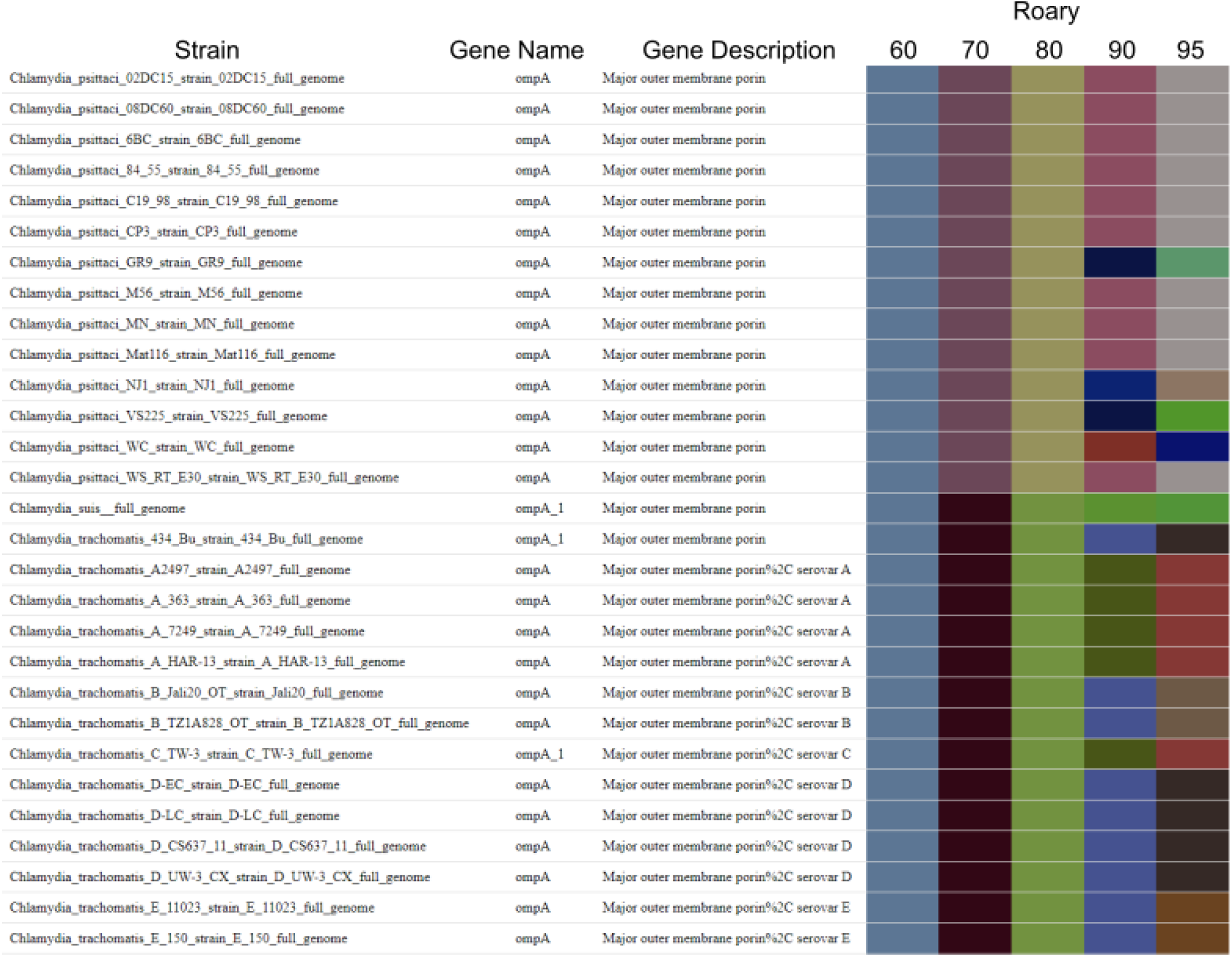
Screenshot of RIBAP HTML output for the *Chlamydia* dataset. For each RIBAP group, the member strains with their annotated gene names and descriptions based on Prokka are presented. Additionally, the user can estimate the sequence similarities of involved genes based on the heatmap representing the individual Roary groups with different sequence similarity thresholds. Here, we selected *ompA*, a gene present in all species of the *Chlamydia* genus. As the colors indicate, Roary failed to sort all *ompA* genes into one cluster with a sequence similarity threshold above 60%. However, its authors do not recommend lowering the sequence similarity threshold to this value (see Roary online FAQ). Further, the HTML output includes a phylogenetic tree for each RIBAP group (not shown in this snapshot) and points to the underlying MSA and NEWICK format file.

## Limitations

There are several important points to consider when using RIBAP for analyzing bacterial genomes. Firstly, when examining larger datasets with more than 100 genomes, the computational runtime and required disk space can become very demanding due to the pairwise gene comparisons and subsequent ILP solving. For example, an input of 32 *Chlamydia* genomes (∼1 Mbp genome size) runs ∼3 h on 8 cores and requires ∼84 GB disk space when using the optional --keepILPs parameter. The disk space can be reduced to ∼2 GB when not storing the intermediate ILP results (default behavior). Running the same dataset on an HPC with the pre-configured SLURM profile reduces the runtime to 1 h. The *Brucella* dataset comprising 71 genomes (∼3.4 Mpb genome size) runs ∼5 h 20 m on an HPC (SLURM default profile) and requires 3.4 TB disk space when keeping the intermediate ILP results. When running in default mode and not keeping the ILPs, the disk space is reduced to ∼16 GB. The results folder has 7.7 GB in both cases. Therefore, we strongly recommend running RIBAP in default mode without saving the intermediate ILP results unless they are really needed for additional examinations. Secondly, while RIBAP performs well on diverse species inputs, it is not as effective when analyzing genomes from the same species. Other established tools, such as Panaroo (Tonkin-Hill et al. 2020) or PPanGGOLiN (Gautreau et al. 2020), predict sound core genomes for intra-species genomes much faster than RIBAP. Thirdly, at the moment, RIBAP does not provide detailed output for core and accessory genomes or persistent/shell/cloud categories as known from other tools. Therefore, RIBAP is most useful for obtaining an accurate estimation of the core gene set for diverse species inputs. Additional metrics have to be extracted from the tabular output RIBAP produces. Further, RIBAP may struggle when analyzing highly similar genes present in multiple copies, such as polymorphic membrane proteins in *Chlamydia*, or genomic regions with high plasticity.

Furthermore, our extension of the proposed ILP is rather simple. Replacing our model with more sophisticated approaches might improve the results of RIBAP further. Recently, Bohnenkämper *et al*. (Bohnenkämper et al. 2021) proposed an extension of the original ILP by Shao *et al*. (Shao, Lin, and Moret 2015) that enables rearrangement analysis of genomes without imposing further restrictions. Expanding from this, Rubert *et al*. (Rubert, Martinez, and Braga 2021; Rubert and Braga 2022) further adapted this model to allow gene family-free analysis of pairwise genomes. Our analysis did not seem limited by the naive ILP model involved. However, future investigations will have to address the question of whether the accuracy of RIBAP can be improved by employing different models to deal with gene duplications and indel events.

Finally, RIBAP, in its current implementation, is also very strict with categorizing genes into the core genome, namely those that are present in all input genomes. Given input data of even higher diversity than in the present study, this conservative threshold could be lowered to, e.g., 95%, which is a generally accepted threshold in other studies as well (called soft core) (Blaustein et al. 2019; Halachev, Loman, and Pallen 2011). In this context, we would also like to point out that better results are possible with the other tools in our comparison by optimizing individual parameter settings. For a fair comparison, however, we have run all tools in their default mode.

## Conclusion

Current computational approaches for calculating the core- and pangenome of diverse input genomes are challenged by low sequence similarities of homologous genes. Therefore, tools tend to underestimate the number of genes present in the core genome of inter-species genomes. Here, we described RIBAP, a pangenome calculation pipeline to overcome this limitation and provide an easy-to-use framework for scientists to analyze pangenomes of diverse input sets. We demonstrated its application to four different bacterial clades and showed the advantage of using RIBAP when genomes from different species of the same genus were the input. Researchers can work exploratively with the RIBAP data and search for genes of interest. The data provided in the HTML report can be used to analyze the presence/absence and sequence diversity within a species or across the species of the genus. However, analyzing core- and pangenomes of bacteria from the same taxonomic clade is not the only use case we envision for RIBAP and pangenomics in general.

Due to the improved detection of gene clusters with low sequence similarity, we see a future application of RIBAP in the study of pan- or core-metagenomes (Ma, France, and Ravel 2020) and the definition of gene clusters in a metagenomic context (Vanni et al. 2022). Determining a core gene set within or between species of metagenomes is highly complicated due to the different species composition and evolutionary distance between bacteria in an environmental sample. However, the principles behind RIBAP are promising to test the application of the pipeline also on metagenome-assembled genomes (MAGs). Thus, high-quality MAGs with high completeness and low contamination could be directly used by RIBAP to identify core genes that shape a comprehensive representation of the genetic content of a taxonomic group in a particular environment.

## Author contributions

KL and MH implemented the pipeline. KL adapted the ILP formulations. KL, LMB, KS, and MH contributed to data analysis and interpretation. KL and MH wrote the first draft of the manuscript. MH conceived the research idea. All authors read and approved the final version of the manuscript and agreed to submit it to the journal.

